# Pre-training artificial neural networks with spontaneous retinal activity improves motion prediction in natural scenes

**DOI:** 10.1101/2024.06.15.599143

**Authors:** Lilly May, Alice Dauphin, Julijana Gjorgjieva

## Abstract

The ability to process visual stimuli rich with motion represents an essential skill for animal survival and is largely already present at the onset of vision. Although the exact mechanisms underlying its maturation remain elusive, spontaneous activity patterns in the retina, known as retinal waves, have been shown to contribute to this developmental process. Retinal waves exhibit complex spatio-temporal statistics and contribute to the establishment of circuit connectivity and function in the visual system, including the formation of retinotopic maps and the refinement of receptive fields in downstream areas such as the thalamus and visual cortex. Recent work in mice has shown that retinal waves have statistical features matching those of natural visual stimuli, such as optic flow, suggesting that they could prime the visual system for motion processing upon vision onset. Motivated by these findings, we examined whether artificial neural network (ANN) models trained on natural movies show improved performance if pre-trained with retinal waves. We employed the spatio-temporally complex task of next-frame prediction, in which the ANN was trained to predict the next frame based on preceding input frames of a movie. We found that pre-training ANNs with retinal waves enhances the processing of real-world visual stimuli and accelerates learning. Strikingly, when we merely replaced the initial training epochs on naturalistic stimuli with retinal waves, keeping the total training time the same, we still found that an ANN trained on retinal waves temporarily outperforms one trained solely on natural movies. Similar to observations made in biological systems, we also found that pre-training with spontaneous activity refines the receptive field of ANN neurons. Overall, our work sheds light on the functional role of spatio-temporally patterned spontaneous activity in the processing of motion in natural scenes, suggesting it acts as a training signal to prepare the developing visual system for adult visual processing.

**Author summary:** Before the onset of vision, the retina generates its own spontaneous activity, referred to as retinal waves. This activity is crucial for establishing neural connections and, hence, ensuring the proper functionality of the visual system. Recent research has shown that retinal waves exhibit statistical properties similar to those of natural visual stimuli, such as the optic flow of objects in the environment during forward motion. We investigate whether retinal waves can prepare the visual system for motion processing by pre-training artificial neural network (ANN) models with retinal waves. We tested the ANNs on next-frame prediction tasks, where the model predicts the next frame of a video based on previous frames. Our results showed that ANNs pre-trained with retinal waves exhibit faster learning on movies featuring naturalistic stimuli. Additionally, pre-training with retinal waves refined the receptive fields of ANN neurons, similar to processes seen in biological systems. Our work highlights the importance of spatio-temporally patterned spontaneous activity in preparing the visual system for motion processing in natural scenes.

## 1 Introduction

Many sensory systems, including the visual system, mature to a large extent prior to the onset of sensory experience. While many different mechanisms support this process, from axon guidance molecules to genetic programs, neural activity is key for the ultimate refinement of synaptic connections from the sensory periphery to downstream areas [1–5]. In the absence of sensory drive, many neural circuits are capable of generating spontaneous neural activity. In the visual system, spontaneous activity is first generated in the retina, where it propagates in wave-like spatio-temporal patterns and is hence referred to as retinal waves [3–5]. Retinal waves have been shown to play a pivotal role in multiple aspects of visual development, including the formation of the retinotopic map [6–8], the refinement of visual receptive fields [6, 9–14], the facilitation of eye-specific and ON-OFF segregation [4, 6, 8, 15], and even the subcellular organization of synaptic inputs [16] in circuits downstream from the retina such as the superior colliculus, the thalamus, and the primary visual cortex. However, the exact mechanisms by which retinal waves contribute to the development of higher-order properties of the visual system, including those for motion processing such as the direction selectivity of neurons found in the superior colliculus and the visual cortex [17–19], remain elusive.

In rodents, retinal waves occur during the first two weeks of postnatal development, while the eyes are still closed, and are classified into three distinct stages with unique cellular mechanisms and propagation characteristics [1, 20]. Recent work showed that retinal waves occurring between the first and second postnatal week exhibit statistical features similar to those observed during visual experience in dynamic environments. Specifically, they display a propagation bias from the temporal to nasal part of the retina [5, 21], mimicking the visual flow that an animal would experience when moving forward through space [21]. Inhibiting this propagation bias resulted in a diminished direction selectivity of downstream neurons in the superior colliculus [21].

Significant theoretical work has investigated the instructive role of retinal waves in the refinement of receptive fields [16, 22], the establishment of long-range connectivity [23], and the retinotopic organization of visual features [9, 15, 24–26] in biologically-inspired networks (see [2, 27] for reviews). However, such networks have not been designed to explicitly probe performance in naturalistic visual computations such as motion processing. On the other hand, recent advances in training artificial neural networks (ANNs) on large samples of visual stimuli have achieved impressive performance, even in the most advanced tasks such as image recognition and segmentation [28, 29]. The structure of the brain historically inspired the development of convolutional and recurrent neural networks, for instance with simple cells providing intuition for the principle of convolutional layers in ANNs [30]. Additionally, studies have shown that the representation of visual information in a convolutional neural network resembles that in the ventral stream of animals [31–35]. Specifically, the processing hierarchy in ANNs, where the layer hierarchy encodes increasingly complex stimulus features, matches that in the animal brain, where complexity increases in downstream brain areas [36–39]. Additionally, spontaneous activity seems to instruct the formation of independent parallel modules in the visual system [40], similar to the modules employed in ANNs. Overall, this substantial line of work speaks to a degree of biological plausibility in ANNs, making them suitable models for studying visual processing.

Previous work has found that pre-training ANNs with retinal waves can improve the downstream performance in image classification tasks and enhance the separability of internal representations learned for each class [41–43]. However, by focusing on image classification tasks, these studies do not consider the temporal evolution of visual input and hence overlook the unique propagation bias of retinal waves.

This prompts a more in-depth investigation of the relationship between the processing of natural visual stimuli rich in motion and the directionally biased propagation of retinal waves. First, we sought to explore whether including biological realism, namely retinal waves as a pre-training signal, would improve the performance of ANNs trained for a naturalistic visual task. Second, we investigated the influence of wave directionality as a spatio-temporally structured feature inherent to spontaneous activity during a particular developmental window. To evaluate the performance of the trained ANNs, we selected a task rich in motion and optic flow, specifically a next-frame prediction task [44], which required the ANN to learn the underlying principles of motion processing. Training and testing ANNs on a natural stimuli dataset mimicking the visual experience of a mouse running through a corridor revealed that ANN performance depended on the training conditions: no pre-training, pre-training with directionally biased retinal waves, and pre-training with retinal waves propagating in random directions. We found that pre-training with retinal waves accelerates learning on natural stimuli, generalizing across ANN pre-training strategies, ANN architectures, and natural stimuli training datasets on which the performance was evaluated. Moreover, we found that pre-training with spontaneous activity refines the receptive field properties of ANN neurons, analogous to observations *in vivo* [6]. Hence, our work suggests that the specific spatio-temporal correlations found in retinal waves are not merely incidental, but are structured to support the development of visual processing of motion in natural scenes in both biological and artificial neural networks.

## 2 Results

### 2.1 Pre-training ANNs with retinal waves accelerates learning for next-frame prediction on natural images

We sought to investigate the effect of pre-training ANNs with spontaneous retinal activity on the ability to process videos with the temporal structure of natural visual stimuli (Fig 1A, B). To achieve this, we trained ANNs to perform next-frame prediction, a task that requires the network to learn both spatial and temporal features of visual inputs. Specifically, we provided 10 consecutive frames of a short movie as input to a three-layer convolutional recurrent neural network trained to predict the subsequent frame. The ANN architecture was designed to mimic the three-tier hierarchical structure of the biological visual system, consisting of retina, thalamus, and primary visual cortex (Fig 1C). The movie either consisted of propagating retinal waves, used as a pre-training signal, or a natural scene with prominent optic flow, employed for training the ANNs (Fig 1D). For the natural scene, we synthesized a dataset that mimics the visual experience of a mouse navigating through a corridor with visual cues on the walls (Methods 4.1.4), a commonly employed paradigm in studies investigating visually guided behavior [45, 46] (Fig 1B). As a second natural dataset, we used the publicly available *CatCam* dataset, which comprises videos recorded in a natural outside setting from a cat’s perspective [47] (Fig 3A).

**Fig 1.**
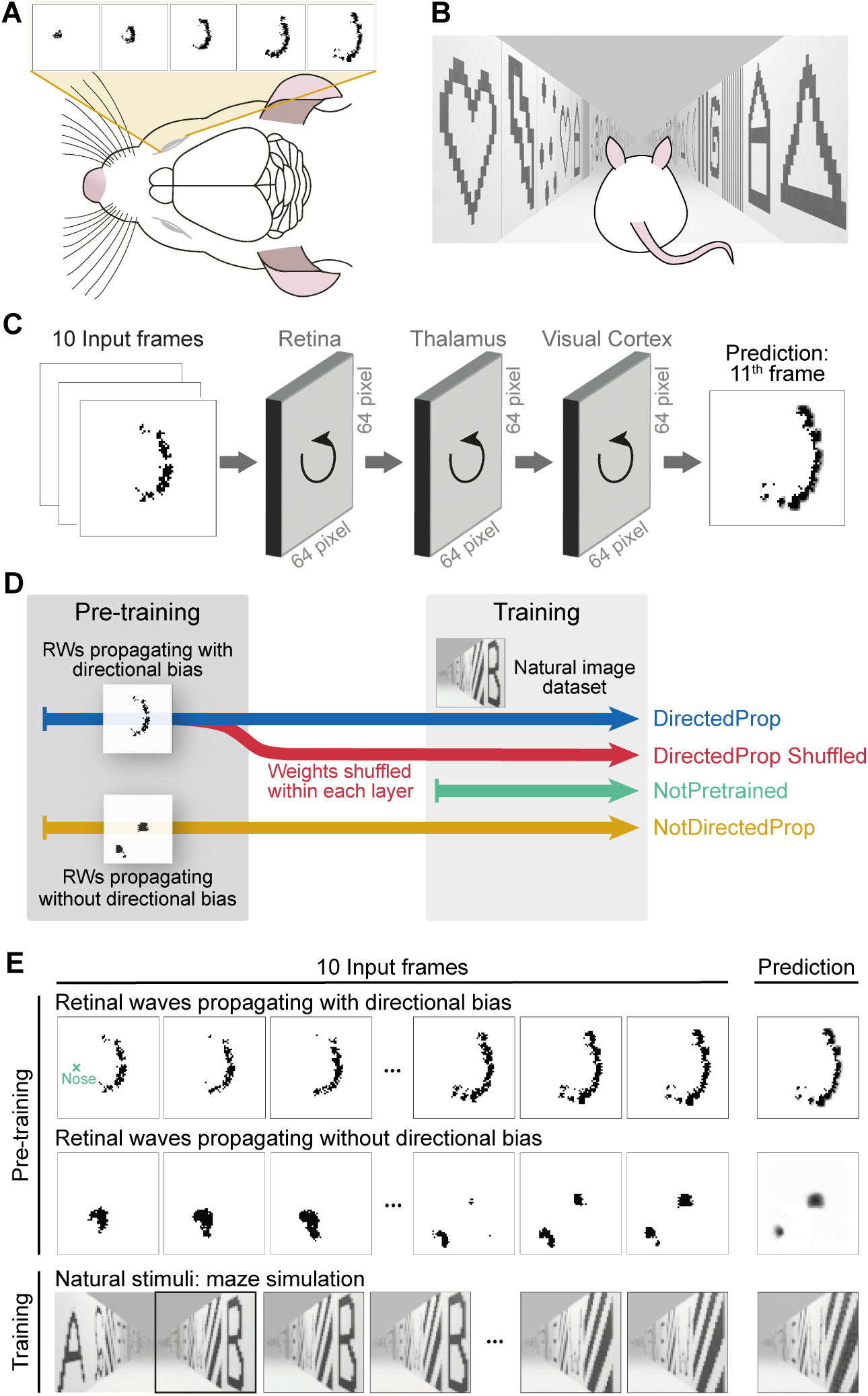
Artificial neural network architecture and training strategy. **A.** A schematic of a developing mouse retina exhibiting retinal waves. Before eye-opening, retinal waves in mice propagate with a directional bias in the animal’s retina during postnatal days 8 to 11. The retinal waves were simulated using a previously published model [24] (Methods 4.1.1). **B.** A schematic of the stimuli seen by the mouse after vision onset in a virtual corridor rich with optic flow, created using a 3D animation software and used as the training dataset (Methods 4.1.4). The dataset simulates a mouse’s visual experience when moving forward through space. **C.** Network architecture. ANNs were trained for the task of next-frame prediction. A sequence of 10 consecutive frames from a video was fed into a three-layer convolutional recurrent neural network, which is trained to predict the 11th frame. **D.** Model training strategies. In total, the performance of four different models was compared: (1) Blue, *DirectedProp* model: Pre-trained with images of retinal waves propagating with a directional bias (Methods 4.1.1), then trained on natural images (Methods 4.1.4). (2) Red, *DirectedProp Shuffled* model: After pre-trained on the dataset comprising retinal waves with directional bias, the weights learned during pre-training were shuffled within each layer before the model was trained on natural images. (3) Green, *NotPretrained* model: Randomly initialized without any pre-training. (4) Yellow, *NotDirectedProp* model: Pre-trained utilizing images of retinal waves without a directional bias (Methods 4.1.2), then trained on natural videos. **E.** Three datasets were used for (pre-)training. The models were pre-trained on either images of simulated retinal waves propagating with a directional bias (top, Methods 4.1.1) or synthetic retinal waves without directional propagation bias (middle, Methods 4.1.2). All models were trained on the dataset depicting a virtual corridor (bottom, Methods 4.1.4), mimicking the right visual field of a mouse upon eye-opening and forward movement.

To evaluate the influence of pre-training using spontaneous retinal activity, we examined the learning trajectories on natural stimuli (corridor simulation or *CatCam* dataset) of four distinct models (Fig 1D,E):

1. *DirectedProp*: pre-trained using simulated retinal waves propagating with a directional bias (Methods 4.1.1), observed at later developmental stages in biological systems (postnatal days 8-9 in mice) [21, 24].
2. *NotDirectedProp*: pre-trained using simulated retinal waves with no directional bias but instead displaying temporal variations in wave size (Methods 4.1.2), observed at other developmental stages [26].
3. *DirectedProp Shuffled*: weights were shuffled within each layer of the *DirectedProp* model after the pre-training stage to test for effects of the weight distribution learned during pre-training.
4. *NotPretrained*: randomly initialized network without pre-training.

To evaluate each model’s capacity to predict future visual scenes, we measured performance by the mean squared error (MSE) (Methods 4.2). To ensure accurate performance measurement and to avoid overfitting, we divided all datasets into training, validation, and hold-out test sets. Pre-training and training were stopped as soon as overfitting was detected. To assess the generalizability of our findings across various three-layered ANN architectures, we investigated “vanilla” RNN, long short-term memory (LSTM), and gated recurrent unit (GRU) architectures for each of the four models (Methods 4.2).

In this section, we introduce exposure to retinal waves as a pre-training stage in addition to training on natural stimuli, thereby increasing the overall training time for pre-trained models (Fig 1D). This tests whether exposure to retinal waves provides a performance advantage at the start of training, suggesting a potential benefit for the nervous system in producing retinal waves to reduce learning time at eye opening. In the subsequent results section, we control for total training time by substituting a portion of the training on natural stimuli with retinal waves, evaluating whether retinal waves offer an advantage that cannot be achieved through natural stimuli alone.

Initially, we studied models employing a vanilla RNN architecture. We found that pre-training on retinal waves (*DirectedProp* and *NotDirectedProp*) substantially enhanced the performance of the ANN on the task of predicting subsequent frames of natural videos featuring corridor navigation during the initial training epochs (Fig 2). When evaluating the performance of models on the simulated corridor validation and test datasets before the first training epoch, without any exposure to natural videos, the models pre-trained on retinal waves showed significantly improved performance compared to the *NotPretrained* model (Fig 2A). The improved performance of the pre-trained models was retained after the first training epoch (Fig 2B). After being fully trained on the natural videos, the performance of all investigated models was comparable (Fig 2C). Statistically significant differences in model performance at this stage were not substantial enough to meaningfully impact frame prediction and are therefore negligible for our goal of comparing the overall visual processing abilities of the models, rather than optimizing performance for each one.

**Fig 2.**
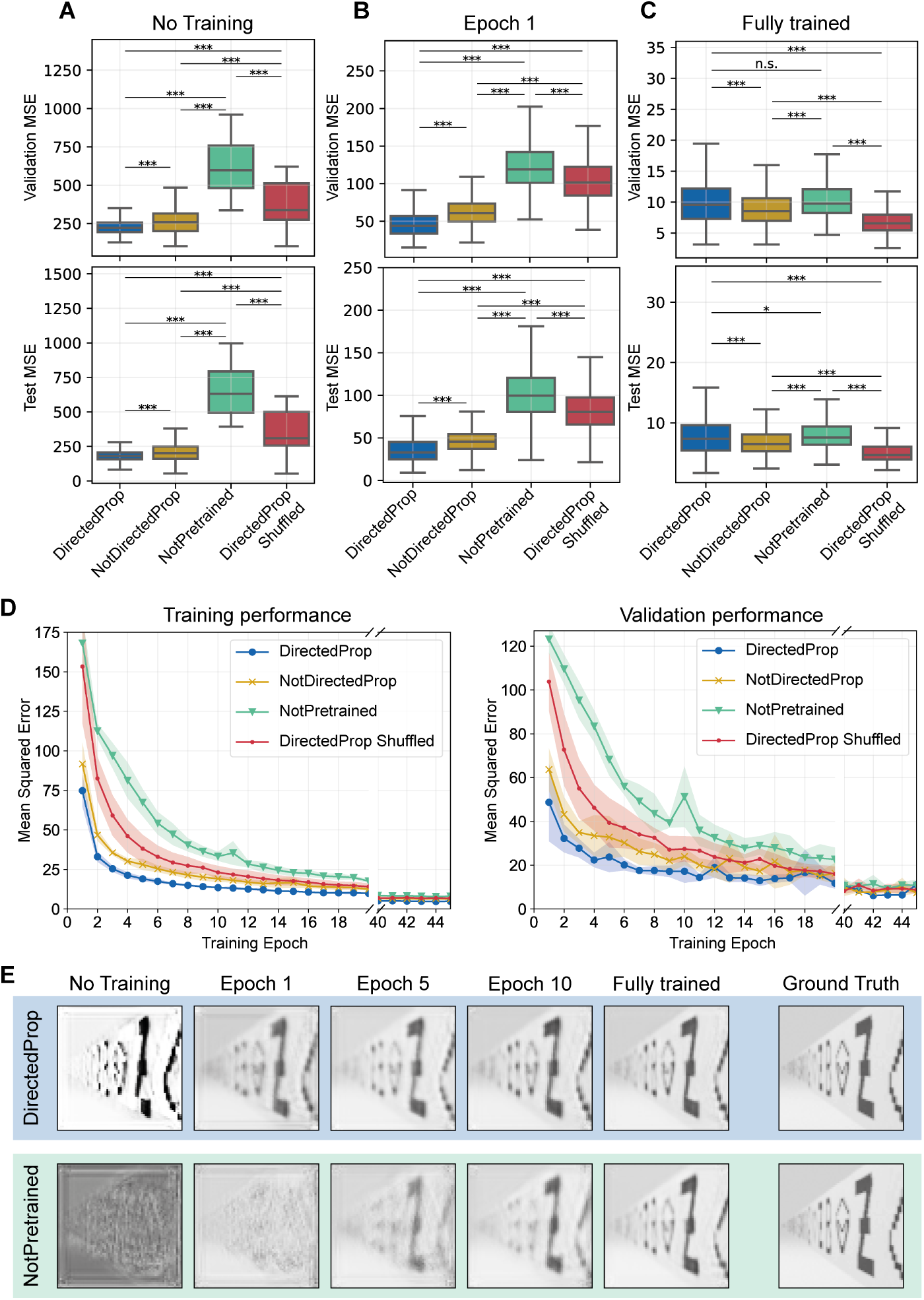
Enhanced performance in next-frame prediction task for natural videos through pre-training with spontaneous retinal activity. **A-C.** Vanilla RNN model performances at different training stages on the simulated corridor validation (top) and test (bottom) datasets. Per model, 5 ANNs were trained using five-fold cross-validation. For the validation dataset, ANN performance was evaluated on the validation set of the respective fold (5 ANNs/folds *×* 200 datapoints per fold = 1000 datapoints in total per model). All ANNs were tested on the same hold-out test set, which comprised 500 datapoints (5 *×* 500 = 2500 datapoints per model). Statistics refer to a two-sided paired t-test (***: *p <*0.001, **: *p <*0.01, n.s.: *p >*0.05), with Bonferroni correction applied. **A.** Performance was evaluated for four models without any exposure to the simulated corridor dataset. **B.** Performance evaluation after one training epoch. **C.** Performance evaluation after training until overfitting was detected. **D.** Learning trajectories of the four vanilla RNN models on the simulated corridor training (left) and validation (right) datasets. Shown is the average MSE across five-fold cross-validation and the corresponding standard deviation. Training continues beyond the epochs shown, with each model stopping individually once overfitting is detected. **E.** Example predictions for the *DirectedProp* (top) and *NotPretrained* (bottom) models at different training stages. The predictions correspond to the 11th frame of a datapoint in the validation dataset. Shown are predictions obtained before any training on the simulated corridor dataset, after one, five, and ten epochs, and following full training until overfitting was detected.

In addition to providing the *DirectedProp* model with a performance head-start on the natural stimuli dataset (Fig 2A), pre-training also accelerates learning thereafter. This became evident when assessing the learning trajectory on natural videos, as the *DirectedProp* model demonstrates a clear learning advantage compared to the *NotPretrained* model (Fig 2D). The finding of accelerated learning is consistent across performance assessments on both the training and the validation datasets (Fig 2D, compare left and right). The accelerated learning in the *DirectedProp* model can be further demonstrated by exploring visual predictions of the models at different training stages. The *DirectedProp* model is capable of predicting key elements of the natural scene, such as the cues displayed on the corridor walls, from early training stages onwards, whereas the *NotPretrained* model requires more training epochs to acquire this ability (Fig 2E).

Both the *DirectedProp* and *NotDirectedProp* models exhibit superior learning compared to the *NotPretrained* model and the *DirectedProp Shuffled* model (Fig 2D). This suggests that pre-training with retinal waves does not merely lead to a different initialization of the weight distribution. Instead, the spatio-temporal features learned from retinal waves during pre-training contribute to the improved performance and are encoded at the level of individual synaptic weights.

To test whether our findings generalize across various three-layered ANN architectures, we further investigated the use of LSTM and GRU architectures as alternatives to the vanilla RNN architecture employed so far. We found that our observations were consistent across all three investigated ANN architectures, despite expected variations in learning speed depending on the complexity of the architecture employed (Fig S1A,B).

To further explore the effect of individual retinal wave characteristics during pre-training on performance with natural videos, we created three additional pre-training datasets. First, to test the impact of wave shape, we generated a dataset featuring propagating squares (Methods 4.1.3), which differ in shape but retain a propagating front. We found that this model performed similarly to the *DirectedProp* model (Fig S2A), suggesting that accelerated learning is not driven by the wave shape but rather by the presence of a clear propagating front. Second, to examine whether sharp black-to-white borders of the propagating wavefront influence performance, we added Gaussian blur to the dataset of retinal waves with a propagation bias (Methods 4.1.1). This model showed substantially accelerated learning and even slightly outperformed the *DirectedProp* model (Fig S2B). This suggests that the network benefits from grayscale colors and blurred borders during pre-training, as these features are also prominent in natural environments. Lastly, to explore the effect of wave directionality, we pre-trained the vanilla RNN architecture with retinal waves that propagate with a directional bias in any direction (contrary to the *NotDirectedProp* model which did not have any bias), as opposed to solely propagating in the temporal-to-nasal direction as observed *in vivo* [21] and used to pre-train the *DirectedProp* model (Methods 4.1.1). Naturally, this dataset also included a fraction of retinal waves that propagate from temporal to nasal. We found that this model performs very similarly to the *NotDirectedProp* model (Fig S2C), corroborating the previous findings: All models pre-trained with spatio-temporal patterns that possess certain characteristics found in spontaneous retinal activity perform substantially better than the *NotPretrained* model during the initial training epochs.

To ensure that any observed performance enhancement is not specific to our simulated corridor dataset, we also tested performance on a dataset featuring prominent optic flow in real natural environments. For this, we used the publicly available *CatCam* dataset of videos recorded in an outdoor environment using a camera attached to a cat’s head [47] (Fig 3A). This dataset is considerably more complex than our simulated corridor videos, as it has three main additional characteristics. Firstly, it incorporates optic flow in all directions, not solely in the direction an animal would experience during forward movement. Secondly, the strength of optic flow varies drastically, ranging from almost no movement to rapid changes due to quick head movements, causing blurry images. Lastly, unforeseen objects can appear in the field of view (Fig 3A). The greater complexity of the *CatCam* dataset also manifests in the deviation of its quantitative characteristics from those in the simulated corridor dataset. Specifically, both the luminance and contrast of images in the *CatCam* dataset show greater standard deviation than in the corridor images (Fig 3B, C), posing a wholly new challenge to the ANNs. Despite the increased complexity, we find that the *DirectedProp* model outperforms the *NotPretrained* model during the initial training epochs (Fig 3D). This finding is consistent across both training and validation datasets. The *DirectedProp* model is capable of predicting key elements of the visual scene, even without any training on natural stimuli (Fig 3E).

**Fig 3.**
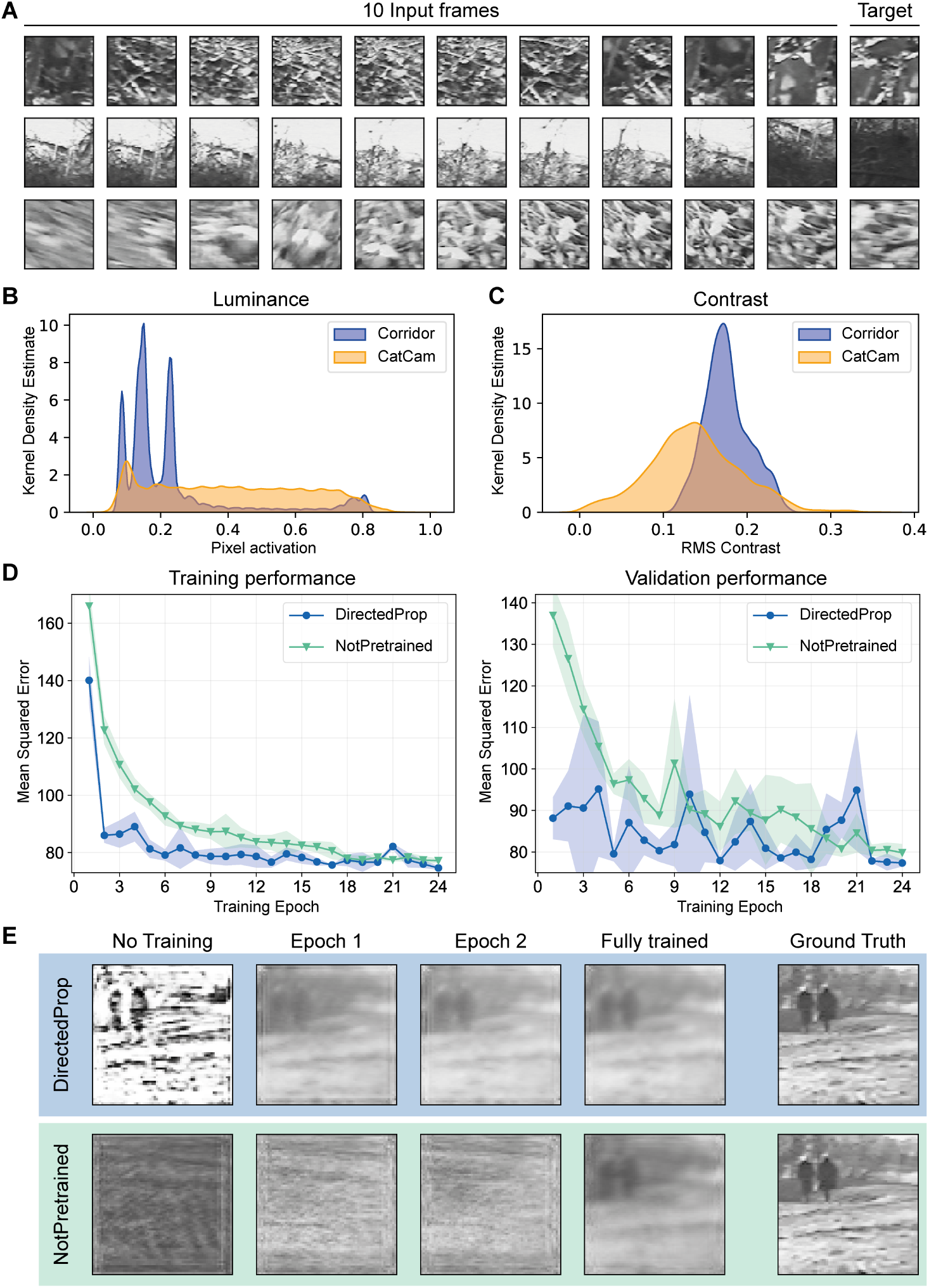
Pre-training with retinal waves improves performance on a complex natural stimuli dataset. **A.** Example datapoints from the *CatCam* dataset. **B.** Comparison of luminance between the simulated corridor (blue) and *CatCam* (yellow) datasets. Luminance was measured as the distribution of pixel activations in the respective training dataset, using the training set of fold 1 for the corridor dataset. Standard deviation: 0.179 (corridor) and 0.222 (*CatCam*). **C.** Comparison of root mean square (RMS) contrast between the simulated corridor (blue) and *CatCam* (yellow) datasets. RMS contrast was computed as the standard deviation of pixel activations for each image in the respective training dataset. Standard deviation: 0.027 (corridor) and 0.054 (*CatCam*). **D.** Learning trajectories of vanilla RNN models on the *CatCam* training (left) and validation (right) datasets. Shown is the average MSE of five replicates per model and the corresponding standard deviation. Training continues beyond the epochs shown, with each model stopping individually once overfitting is detected. **E.** Example predictions for the *DirectedProp* (top) and *NotPretrained* (bottom) models at different training stages. The predictions correspond to the 11th frame of a datapoint in the validation dataset. Shown are predictions obtained before any training on the *CatCam* dataset, after one epoch of training, after two epochs, and following full training until overfitting was detected.

In sum, we found that pre-training with spontaneous retinal activity provides the ANN with a performance head-start when predicting videos featuring natural stimuli, and results in accelerated learning thereafter. This suggests that retinal waves offer a sufficiently rich signal to prepare the developing visual system for real-world motion processing.

### 2.2 Pre-training on retinal waves accelerates learning compared to training exclusively on natural stimuli despite equal training time

Based on the finding that pre-training accelerates learning on the next-frame prediction task for natural videos, our next objective was twofold: first, to exclude that this improvement is solely due to the longer training time resulting from pre-training, and second, to determine how the duration of pre-training influences the observed performance enhancement. To address this, we created models for which we replaced one, two, or four training epochs on the simulated corridor dataset with pre-training on retinal waves, rather than having an additional pre-training stage (Fig 4A). Hence, when comparing performances at a specific training stage, all models had been exposed to an identical number of datapoints, corresponding to one 11-frame video sequence each.

**Fig 4.**
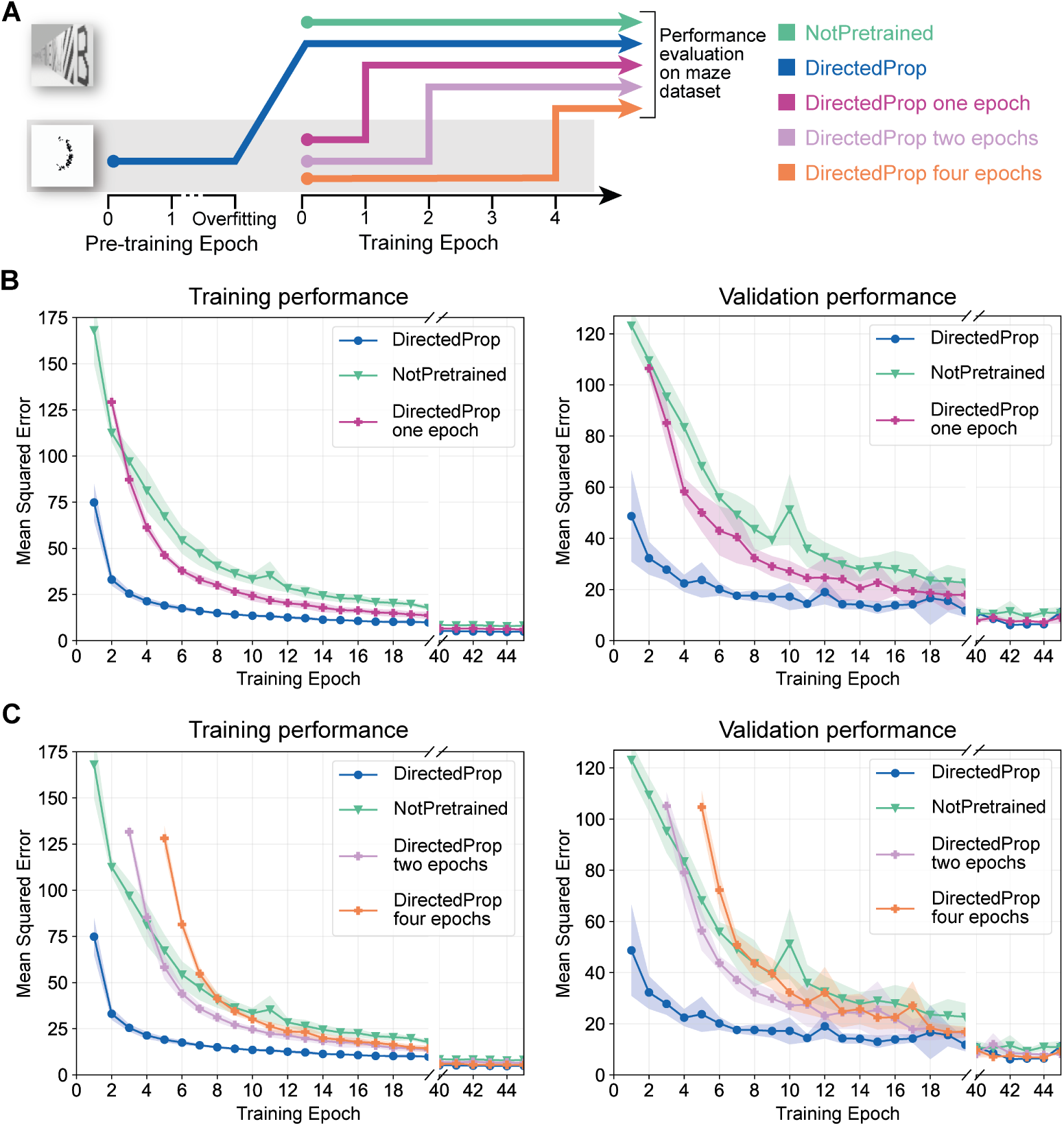
The pre-trained model shows accelerated learning compared to a model trained exclusively on natural videos despite equal training time. **A.** Training strategy for model comparison with equal training time: Instead of incorporating a separate pre-training stage alongside training on natural stimuli, we substitute initial training epochs with pre-training on retinal waves. **B.** Learning trajectories of three vanilla RNN models on the simulated corridor training (top) and validation (bottom) datasets. Shown is the average MSE across five-fold cross-validation and the corresponding standard deviation. Note that the performance assessment for the model pre-trained for one epoch on retinal waves (*DirectedProp one epoch* in pink) starts at epoch two, ensuring direct comparability to the *NotPretrained* model as this model does not have a longer training time. Training continues beyond the epochs shown, with each model stopping individually once overfitting is detected. **C.** Learning trajectories of vanilla RNN models on the simulated corridor training (top) and validation (bottom) datasets. In addition to the *DirectedProp*and the *NotPretrained* models, we evaluated the performance of models for which either two or four training epochs were replaced with pre-training on retinal waves.

For the models with a vanilla RNN architecture, we found that pre-training on retinal waves for only one epoch (*DirectedProp one epoch*) already substantially speeds up learning on the natural video dataset (Fig 4B). Although the model does not achieve as small a loss as the *DirectedProp* model (pre-trained until detection of overfitting) during the initial training epochs, the *DirectedProp one epoch* model still shows accelerated learning compared to the *NotPretrained* model. Consistent with the findings from the previous section, these observations generalize across both training and validation datasets (Fig 4B), as well as across LSTM and GRU architectures (Fig S3). These results are surprising considering the pre-trained model had a shorter training time on the natural video dataset, which excludes the possibility that the accelerated learning observed for pre-trained models is solely attributable to the longer training time. Instead, our results imply that the pre-training on spatio-temporally complex patterned retinal waves uniquely influences the ANN weights in a manner not achievable through immediate exposure to real-world natural stimuli.

We further explored the learning trajectories of vanilla RNN models where either two or four training epochs were replaced with the corresponding number of pre-training epochs on retinal waves (*DirectedProp two*/*four epochs*). For both models, we observed accelerated learning compared to the *NotPretrained* model. The pre-trained models achieve a smaller MSE after 2-4 epochs of training on the natural videos, despite having been trained on fewer natural stimuli datapoints at that point (Fig 4C). Consistent with the findings in the previous section, also the performance enhancement for the models with fewer pre-training was not persistent, resulting in comparable performance among all models after being trained on the simulated corridor dataset until overfitting was detected.

Our simulated corridor dataset includes several effects inherent to natural scenes, such as color shifts (increased contrast toward the front) and size differences (objects appear larger as they move closer to the front), which the network must learn for accurate predictions. To isolate these effects, we created separate training datasets, each focused on one feature: either size or color changes in a gradually transforming circle (Methods 4.1.5). Evaluations on the *DirectedProp*, *NotPretrained*, and *DirectedProp one epoch* models showed that both the *DirectedProp* and *DirectedProp one epoch* models show accelerated learning compared to the *NotPretrained* model for predicting color (Fig 5A,B) and size (Fig 5C,D). These findings, consistent with those from the corridor dataset, suggests that pre-training with retinal waves, or even just replacing the first training epoch with retinal waves, prepares the ANN to subsequently learn and predict various environmental features. This includes changes in color and size, both of which are essential for navigating natural stimulus environments.

**Fig 5.**
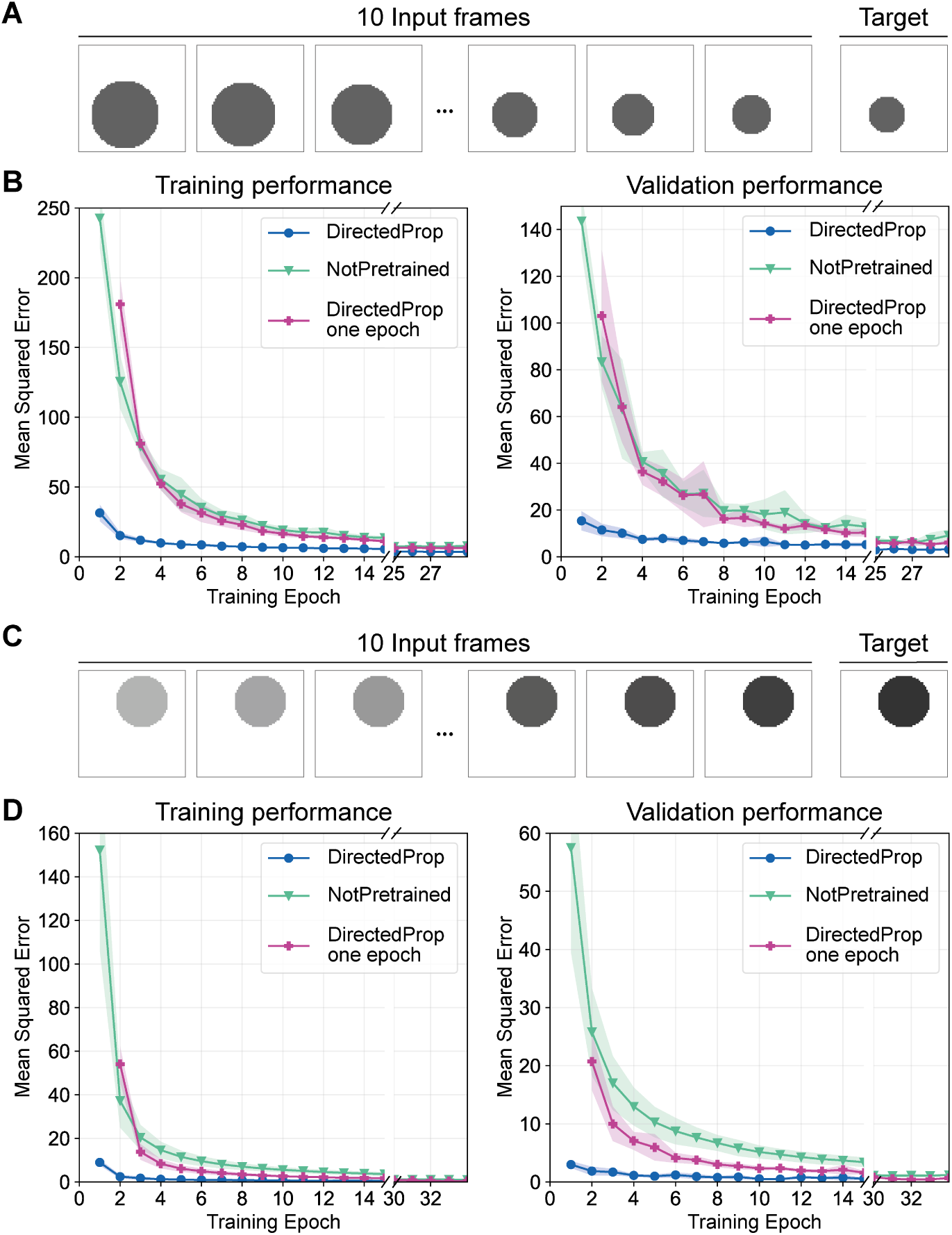
Pre-training with retinal waves accelerates learning of both color and size changes. **A.** Example datapoint from the training dataset comprising circles with gradually changing size (Methods 4.1.5). **B.** Learning trajectories of four vanilla RNN models on the size-progression training (left) and validation (right) datasets. Shown is the average MSE across five-fold cross-validation and the corresponding standard deviation. Training continues beyond the epochs shown, with each model stopping individually once overfitting is detected. **D.** Same as panel A, but for the dataset comprising circles with gradually changing color (Methods 4.1.5). **E.** Same as panel B, but for the color-progression dataset.

Overall, the transient acceleration in the learning process, which is not simply due to the longer training time for the task, suggests an important role of retinal waves in setting up the weights in the network. The findings indicate that spontaneous, spatio-temporally patterned activity in the retina prior to the onset of vision is capable of priming the visual system in a manner that direct exposure to natural stimuli cannot achieve.

### 2.3 Pre-training refines the spatio-temporal receptive field of ANN neurons

To explore ANN characteristics underlying the improved performance resulting from pre-training with spontaneous retinal activity, we investigated the receptive field properties of the models (Methods 4.3). Given that retinal waves are known to refine the receptive fields of neurons in the developing visual system *in vivo* [6, 9–14], we asked whether a similar effect could be observed in ANNs. Specifically, we compared the properties of the receptive field of one target neuron in the output layer across the different training conditions (Fig 6A). To extract receptive fields, we employed an activation maximization algorithm [48], which uses gradient ascent to find the inputs to the ANN that maximally activate the target neuron in the output layer after processing all the input frames (Methods 4.3). Given the nature of our next-frame prediction task, we investigated the receptive field properties in both in the spatial and temporal dimensions. Based on our results thus far, we restricted our exploration of receptive field properties to the vanilla RNN architecture.

**Fig 6.**
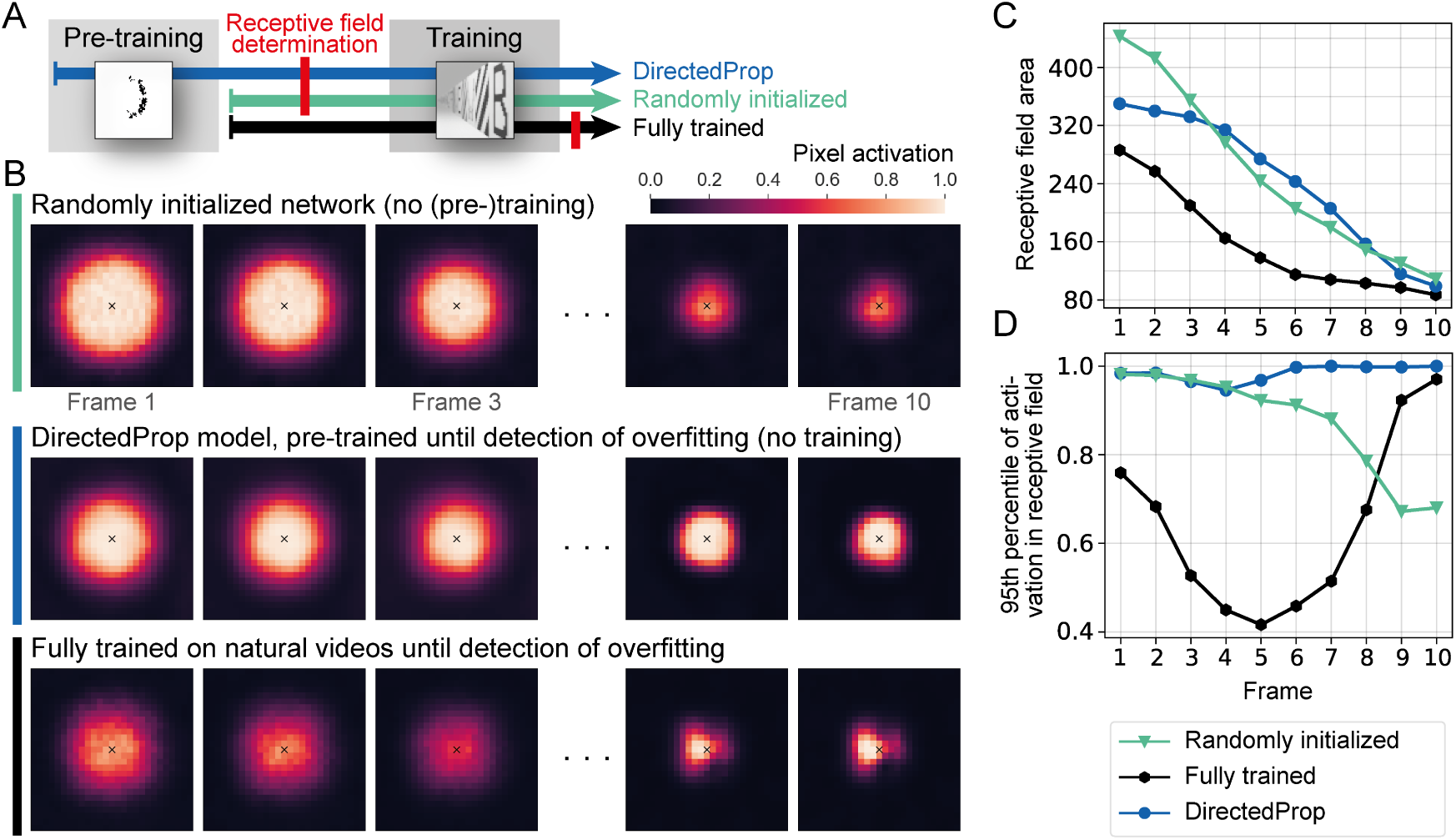
Pre-training refines the spatio-temporal receptive field of neurons in the output layer. **A.** Receptive fields were determined for three models, all using a vanilla RNN architecture: a randomly initialized model without any training prior to investigating the receptive field properties (green), a model trained on the natural videos until detection of overfitting (black), and a model pre-trained on retinal waves with directional bias until detection of overfitting, without any training on natural videos (blue). **B.** Spatio-temporal receptive field of a target neuron in the last 2D convolutional layer of the ANN. Shown are the 10 input frames, each cropped to a square of 27 *×* 27 pixels, which maximally activate the target neuron, indicated by a cross. The displayed result represents the average of 50 trials (10 trials for each of the 5 ANNs per model) with 100 optimization steps per trial. Notably, the decrease in the receptive field size over the 10 frames for each model is a result of the convolution process. **C.** Receptive field area for each frame, defined as the number of activated pixels, of the models described in A. Shown are the properties of the receptive fields averaged over the 50 trials per model, as described in B. **D.** Receptive field strength, measured as the 95th percentile of pixel activations, for each frame.

We first analyzed the receptive field size, measured by the area in pixels, and found that, over the 10 input frames, the receptive field size decreases in all training conditions (Fig 6B). Given that successive convolutions increase the receptive field size in the input space, the last input frame, which is processed only through the feedforward pathway, is expected to have a smaller receptive field than the first input frame, which mainly influences the computation via the recurrent connections. For the first input frame, both pre-training on retinal waves and training on natural videos result in a decrease in receptive field size compared to a randomly initialized ANN (Fig 6B,C). This suggests that (pre-)training with retinal waves refines the receptive field to be more local, bringing its properties closer to what is found after training on natural videos.

In addition to differences in receptive field size, the receptive field strength, which indicates the contribution of the respective frame to the activation of the target neuron, also undergoes changes during (pre-)training. The model trained fully on natural videos demonstrates a reduced receptive field strength in the first 8 input frames when compared to the randomly initialized model (Fig 6D). However, for frames 9 and 10, the strength increases for the model trained fully on natural videos (+37% for frame 10). Likewise, for the model pre-trained with retinal waves (*DirectedProp* model), the strength in the last frame is increased (+40% for frame 10). This suggests that (pre-)training directs the neuron’s attention more towards the last input frames rather than the earlier ones. Considering the next-frame prediction task, this appears to be a valuable ANN characteristic, as input closer in time to the desired prediction holds more information for the task compared to earlier time frames.

Only marginal differences were observed across models with different pre-training strategies, namely the *NotDirectedProp* model, the *DirectedProp* model pre-trained for one epoch, and the *DirectedProp* model pre-trained until detection of overfitting (Fig S4). All three models demonstrate a decreased receptive field size for the initial input frames and an increased strength for the final input frames compared to a randomly initialized network. This aligns with the finding of similar learning trajectories for all models pre-trained on retinal waves (Fig 2D and Fig 4B,C).

These results highlight the effectiveness of pre-training with retinal waves in refining the spatio-temporal receptive field properties of ANNs. Specifically, pre-training leads to a decrease in size in the first input frames and an increase in strength in the last input frames, bringing the receptive field properties closer to the characteristics achieved through training on natural stimuli. This refinement of receptive fields, aligning with observations made *in vivo* [6], could represent one potential mechanism explaining the accelerated learning of pre-trained models on natural videos.

### 2.4 Pre-training refines the neuronal response variability

To further investigate the effect of pre-training with retinal waves on ANNs, we conducted a comparative analysis of the neuronal response characteristics to grating stimuli across different models (Methods 4.4). We designed 120 grating stimuli based on the Allen Brain Observatory: Visual Coding dataset [49], originally developed to characterize the tuning properties of visual cortical neurons. As in previous analyses, we focused on the vanilla RNN architecture, consisting of three convolutional recurrent layers and one output layer. Neuronal responses for the different models were assessed at the same training stages as in the receptive field investigation (Fig 6A). To evaluate neuronal response properties, we compared the activity of a neuron when presented with a grating stimulus (Fig 7A) to its activity when exposed to a full-field baseline stimulus of identical mean luminance (Fig 7B). The distribution of activity changes between each of the 120 grating stimuli and the corresponding full-field stimulus revealed differences between the models when examined at the level of individual neurons. We found that in all ANN layers, pre-training with retinal waves shifted the neuronal response variability very close to that of the model trained on natural stimuli (Fig 7C,D), consistent with our previous findings on the receptive field properties of ANN neurons. While the neuronal response variability was reduced in the three recurrent layers for the DirectedProp and fully trained models compared to the randomly initialized network, the variability of neuronal responses in the output layer increased (Fig 7C,D). A higher neuronal response variability in the output layer indicates that the predicted frames differ depending on whether the ANN was presented with a grating or a full-field stimulus. This suggests that exposing the ANN to spontaneous retinal activity or natural stimuli improves the network’s ability to process and respond to the input, providing an additional explanation for the accelerated learning observed in the DirectedProp model.

**Fig 7.**
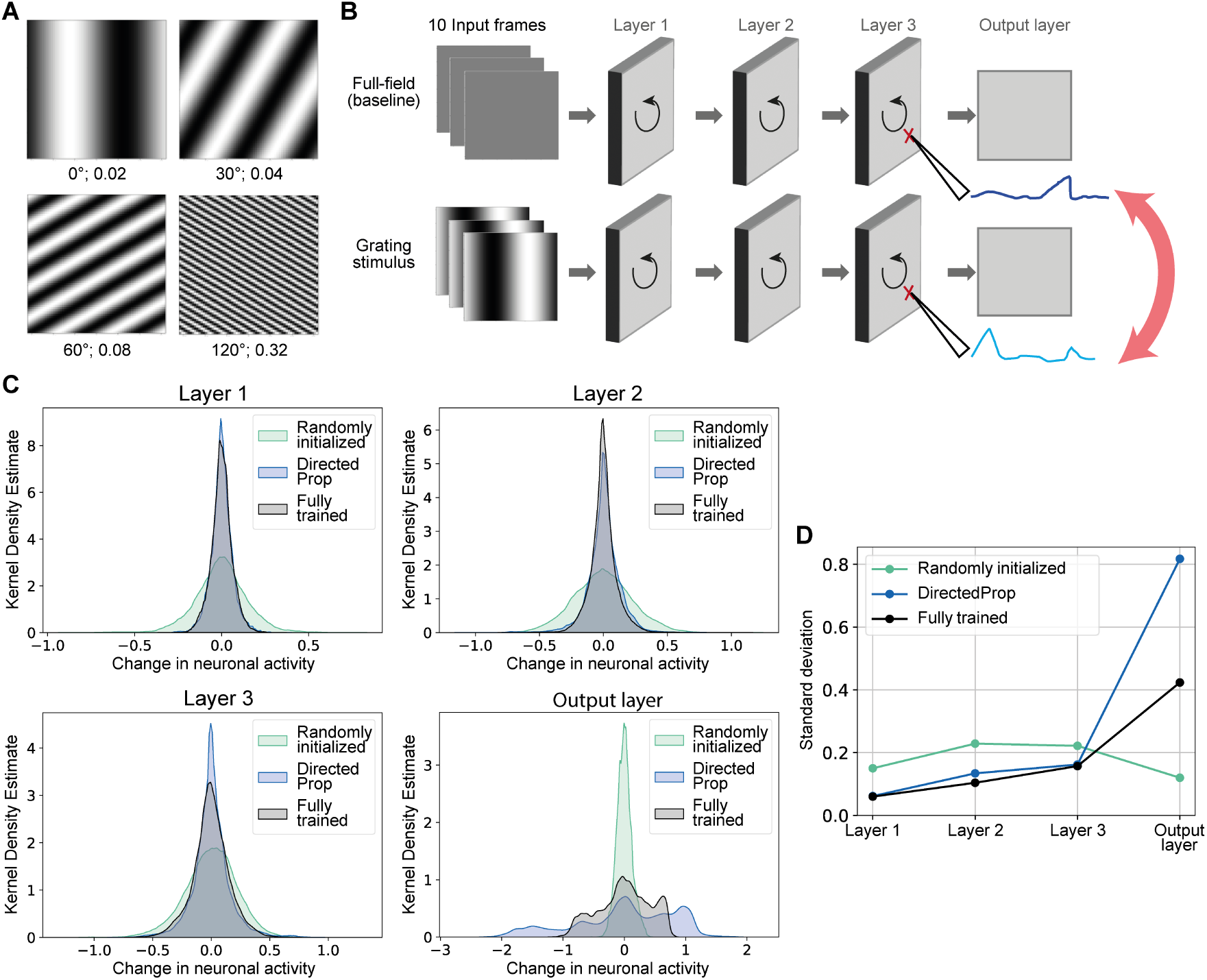
Pre-training refines the neuronal response variability. **A.** Examples of static grating stimuli with varied orientations and spatial frequencies. **B.** Schematic of the static grating experiment. A grating stimulus with a distinct orientation and spatial frequency was paired with its corresponding blank stimulus, a uniform gray field matching the mean luminance. Both stimuli were presented for 10 frames, with the average neuronal response computed over this duration. The mean response to the blank stimulus served as a baseline for calculating the change in activity for each neuron (Methods 4.4). **C.** Kernel density estimation of the distribution of neuronal response changes across all neurons and all grating stimuli within a given layer for each of the three models investigated (120 *×* 64 *×* 64 *×* 64 datapoints for each hidden layer and 120 *×* 64 *×* 64 datapoints for the output layer). **D.** Standard deviations of the various distributions of neuronal response changes shown in panel C.

## 3 Discussion

Spontaneous neural activity before the onset of sensory experience is observed in many neural circuits during development and plays an instructive role in the formation and refinement of neural connections [1, 50–52]. Here, we focus on spontaneous activity in the developing visual system, specifically in the retina, which has been shown to induce refinement of connections in downstream visual areas such as the thalamus and visual cortex [1, 4].

Using ANNs, we demonstrated that retinal waves could play an instructive role in priming the visual system for motion processing upon eye-opening. We found that pre-training neural networks with retinal waves temporarily improved performance on a visual task rich in motion, even when the total training time was matched by simply replacing initial training with pre-training. The observed performance advantage is consistent across several ANN architectures (vanilla RNN, LSTM, GRU) and across natural movie datasets containing both simulated and real-world examples of optic flow. In a biological context, this enhancement would correspond to an improved motion processing capacity immediately upon eye-opening. Importantly, our results suggest that retinal waves prime the visual system in a way that direct visual experience cannot achieve. A possible explanation for this could be that the statistical simplicity of retinal waves is more effective in setting up network properties, such as receptive fields and neuronal response variability, than complex natural stimuli. Supporting the importance of low-level stimuli at early developmental stages is the finding that ANNs show improved information integration across larger image areas and better face recognition capability across a range of image resolutions when trained on a sequence of blurry to high-resolution images rather than on high-resolution face images only [53].

We explored the impact of pre-training with retinal waves reflective of different developmental stages: those with a directional propagation bias (postnatal days 8 to 11 in mice), and those merely increasing and decreasing in size with no propagation bias. We observed a meaningful improvement in performance during the initial training epochs for all retinal wave varieties. We noted a slight performance improvement when using retinal waves with a temporal-to-nasal bias; however, this advantage was minimal compared to the performance difference observed in comparison to the model without any exposure to retinal waves. This implies that the mere presence of patterned spontaneous activity, rather than the propagation bias during a specific developmental time frame, might be driving enhanced motion processing. Notably, this aligns with the finding that horizontal direction selectivity maps were diminished in mice with reduced retinal wave frequency in the first postnatal week but maintained wave activity with propagation bias in the second week [54]. Hence, in biology, the propagation bias does not seem to drive the establishment of direction selectivity maps [54], and similarly, in our case, it has minimal influence on the ability to accurately predict motion as a higher-level visual task.

Investigating the receptive field and neuronal response properties of our ANN models revealed that pre-training on retinal waves shifts these characteristics toward those observed when training on natural images until overfitting is detected. Specifically, we observed a reduction in receptive field size in the first input frames, and an increase in strength in the last frames. The decreased receptive field size in (pre-)trained models aligns with findings from *in vivo* studies, where disrupted retinal wave activity has been associated with unusually large receptive fields in mice [55–57]. Given that the last input frames are temporally closest to the target prediction and therefore contain information highly relevant to the task, the increased attention of the neuron towards those frames provides one plausible explanation for the improved performance observed for pre-trained models during the initial training on natural stimuli. Furthermore, our analysis of neuronal responses revealed that exposing an ANN to retinal waves and natural videos enhances the differences between frames predicted from grating versus full-field input stimuli, suggesting an improved ability to process and respond to specific sensory stimuli.

Although we investigated artificial neural networks, this work allowed us to gain insights into the functionality of biological visual systems. Nonetheless, several key components of our study setup had limited biological plausibility. (1) While the concept of pre-training ANNs with spontaneous activity has been investigated before [41–43], here we specifically focus on the motion and directionality inherent to retinal waves, and the possible advantage thereby endowed to motion processing in the visual system. To constrain the ANN to learn motion features, we used a task that could only be solved once the ANN had learned the mechanisms underlying motion and optic flow: next-frame prediction. In contrast, previous work used image classification as the evaluation task [41–43], providing only static images and neglecting the temporal dimension of input to the visual system. The ability to process natural stimuli, detect moving and immobile objects, and predict future motion is crucial for animals, for instance in the event of a predator attack. We designed the simulated corridor dataset to encompass these features by incorporating components that appear constant (floor and ceiling), while the side walls depict strong movement. In combination with the next-frame prediction task, this represents a higher-order visual task that includes biologically relevant features.

(2) We verified the reproducibility of our findings across vanilla RNN, LSTM, and GRU architectures, all of which use convolutional kernels to process the provided images. The information processing in a convolutional neural network and the visual system bears remarkable similarities, such as the hierarchical representation of increasingly complex features [36–39] and the formation of independent parallel modules driven by spontaneous activity in development [40]. Despite certain properties being different from the animal brain [58], this suggests some extent of biological plausibility in such architectures.

(3) To train the ANNs in this study, we used backpropagation, an algorithm typically considered incompatible with biological principles [59, 60]. While Hebbian learning offers a more brain-like learning rule [61], implementing it for our supervised task of next-frame prediction has proven challenging. Previous work that explored the impact of retinal waves on ANNs in the context of image classification implemented a Hebbian learning rule only for the pre-training stage [41], which poses a potential future direction for our work. Another algorithm that emerged out of the desire for more biologically plausible learning algorithms is predictive coding, which, unlike backpropagation, assumes learning occurs through the continuous update of internal representations by constant comparisons to the outer world [62, 63]. An interesting continuation of our work would be to explore the effect of pre-training with retinal waves on ANNs designed for next-frame prediction but trained using predictive coding [64], which has been shown to exhibit many characteristics originally observed in the biological visual system [65].

Another important question to consider pertains to the uniqueness of retinal waves as a pre-training signal. While we found that simple synthetic patterns, such as propagating squares, can also lead to performance improvements, this does not diminish the distinct biological relevance of retinal waves. Their significance may lie in their potential generation by minimalistic, genetically encoded circuits [66, 67]. While engineered patterns can produce similar training effects, they may lack the same efficiency of naturally occurring retinal waves in a biological context.

In sum, our computational models enabled us to explore the functional role of spontaneous retinal activity during development in preparing the visual system for higher-order visual tasks. Our results suggest that retinal waves prime the visual system for future motion processing in a way that direct exposure to natural stimuli cannot achieve.

## 4 Methods

### 4.1 Dataset creation

In total, three datasets were generated for this study: two comprised images of retinal waves and were used for pre-training, while the third training dataset consisted of natural videos. Each dataset was divided into training and validation sets, allowing for detection of overfitting and an accurate performance assessment. Both retinal wave datasets utilized for pre-training comprised 2000 datapoints in the training and 300 datapoints in the validation set. For the subsequent training dataset, 500 and 200 datapoints were allocated to the training and validation sets, respectively. When comparing models with equal overall training time (Fig 4), the retinal wave pre-training dataset was downsampled to the same size (500 and 200 datapoints for training and validation set, respectively), to ensure full comparability between the models. Each datapoint was stored as a 11 × 64 × 64 array, where the first 10 frames of size 64 × 64 pixels served as input frames, and the eleventh frame represented the ground truth prediction, corresponding to the label in a supervised machine learning setting.

#### 4.1.1 Pre-training dataset comprising retinal waves with directional bias

For the models pre-trained with retinal waves exhibiting a directional propagation bias, a dataset comprising simulated end-of-stage 2 (P8-P9) retinal waves was used. These waves were generated using a previously published model that simulates the propagation of exactly one wave at any timepoint [24]. The model generates waves in a biologically inspired manner by implementing a source of asymmetric inhibition from which the wave propagates away, as observed *in vivo* [21]. Following the approach outlined in the previously published model, we used a square grid on which the waves propagate, where each position in the grid can be interpreted as representing both an ON and an OFF retinal ganglion cell (RGC). We used a retina/grid model with dimensions of 64 × 64, which also matches the frame size of the generated images, so that one pixel corresponds to one RGC location. To initiate a wave, an initiation position was randomly sampled from a uniform distribution across the grid. Furthermore, a point of asymmetric inhibition was sampled, determining the direction of wave propagation.

In this study, we primarily used a dataset with retinal waves propagating with a temporal-to-nasal bias over the retina (DirectedProp model). However, for comparison, we also briefly explored vanilla RNN models pre-trained with retinal waves featuring directionally biased propagation towards all directions, not just temporal-to-nasal. The only difference between the wave generation for these two datasets is the procedure for sampling the asymmetric inhibition point. For the waves propagating in all directions, this point is randomly sampled from a uniform distribution across the grid. To generate waves with directional bias, the asymmetric inhibition point was sampled from a 2D Gaussian distribution, centered at a position considered as “nose” (pixel [32, 8] in a numpy array representing the grid) with a spread of *σ* = 1 pixel. As proposed by the previously published model, a local propagation bias *b* = 0.35 was used to simulate larger, curved waves, resembling those observed at the end of stage 2 [21]. Although we simulated the retinal waves with a delay in OFF RGC activation [24], the resulting images depicting the propagating waves were subsequently transformed into black and white representation, yielding a binary firing pattern that does not differentiate between ON and OFF RGCs.

Let *F_w_* denote the total number of frames in which the retinal wave *w* propagated, defined as the presence of at least one activated RGC in each frame. Only waves *w* with *F_w_ >* 11 were included in the dataset, as the creation of one datapoint requires the presence of at least 11 consecutive frames. To ensure the ANN has the opportunity to learn wave characteristics at all stages of propagation, a starting frame *t* ∊ [1*, F_w_* – 11] was randomly chosen for each wave *w*. Finally, to generate one datapoint per simulated wave, we stored the frame sequence [*t, t* + 11] as one datapoint, with frame *t* + 11 corresponding to the frame the ANN is trained to predict.

In addition to the binary retinal wave images, where each pixel represents an RGC that is either firing or not, we also explore retinal waves with added Gaussian blur as a pre-training signal. To achieve this, we apply a Gaussian filter (*scipy.ndimage.gaussian filter* with *sigma=0.7*) to each image in the dataset comprising retinal waves propagating with a temporal-to-nasal bias.

#### 4.1.2 Pre-training dataset comprising retinal waves without directional bias

The dataset comprising retinal waves propagating without a directional bias consists of waves generated through a previously published model [26]. This model differs from the one employed for generating directionally biased waves in two ways: Firstly, it generates spontaneous activity that increases and decreases in size rather than propagating from one end of the retina to the other. Secondly, it allows for the presence of multiple or no propagating retinal waves at any given timepoint.

The model includes a simulation of amacrine cell activity, simulated calcium response, and the temporal wave progression. To obtain images only of those cells active at the timepoint of recording, we opted to obtain images depicting the simulated amacrine cell activity from the model. After recording images for a sufficient duration, the images were resized into 64 × 64 frames using exact nearest neighbor interpolation (*opencv2.resize* with *interpolation=cv2.INTER NEAREST EXACT*) and transformed into black-and-white images to match the binary firing pattern in the first pre-training dataset. To ensure effective learning of wave characteristics, we included only frames in which more than 5 cells out of the total number of 64 × 64 = 4096 cells were active in further downstream processing. Finally, datapoints comprising 11 consecutive frames were extracted, with a starting point every four frames.

#### 4.1.3 Pre-training dataset comprising propagating squares

For the pre-training dataset featuring squares moving across the image, we aimed to match the number of activated pixels to that in the retinal wave pre-training dataset. Specifically, for each datapoint in the retinal wave dataset, where waves propagate with a temporal-to-nasal bias, we define *ρ* as the maximum number of RGCs firing (i.e., activated pixels) in any frame across all 11 frames. For the corresponding datapoint in the moving square dataset, we set the square’s side length to 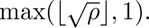 A starting position was uniformly sampled, positioning the square within the frame (i.e., at most touching the border), along with a movement direction randomly chosen from eight possible directions (up, down, left, right, up-left, up-right, down-left, down-right). Over the 11 frames, the square was then moved in the selected direction at a speed of two pixels per frame.

#### 4.1.4 Training dataset featuring a simulated corridor

The training dataset was designed to resemble natural videos with strong optic flow as experienced by an animal when moving forward. We aimed to mimic the visual experience of a mouse navigating through a corridor, a frequently used approach when studying the visual system [45]. For this, we generated visual cues to be projected onto the walls of a digitally animated corridor that extends infinitely in a straight path.

To assess performance on unseen data, we conducted five-fold cross-validation on this dataset. In addition to the training and validation datasets, we created a hold-out test set of 500 datapoints to evaluate performance on data completely independent of the training process. To ensure accurate performance assessment, the cues projected on the walls in the training, validation, and test sets were distinct from each other. Consequently, to enable five-fold cross-validation, we partitioned the wall cues into six disjoint subsets of 14 cues each. Five of these subsets were used to create, respectively, one training dataset with 500 datapoints, and for each training dataset, 200 datapoints from a different fold were used as the validation dataset, ensuring no overlap in wall cues between training and validation. The sixth subset of wall cues was used to generate the entirely independent hold-out test set. To simulate the corridor environment, we leveraged the 3D animation software Blender and custom Python scripting to automatically position the wall cues within the virtual environment. Frames capturing the right visual field were rendered at a resolution of 64 × 64 pixels, and subsequently, sequences of 11 consecutive frames were saved as individual datapoints.

#### 4.1.5 Training datasets comprising looming stimuli

We generated two additional training datasets comprising looming visual stimuli, as used in previous studies of the visual system [68]. Firstly, 11-frame sequences featuring a circle that steadily increased or decreased in size were created. The direction (increasing or decreasing) and the center pixel position (*x* and *y* coordinates, respectively, from [10, 54)) were uniformly sampled. For increasing stimuli, the starting radius was sampled from [5, 20), and for decreasing stimuli, from [20, 40). A random grayscale color (from black to white) was also assigned.

Secondly, in the dataset with circles changing color across frames, the center position, a radius in the range [5, 20), and a direction (brightening or darkening) were sampled. The starting color was randomly sampled from [0, 0.5] for brightening stimuli and [0.5, 1] for darkening stimuli. The color changed incrementally over 11 frames, with a step size of 0.05 (minimum 0, maximum 1).

### 4.2 ANN architectures and training

The 2D convolutional recurrent neural network architectures were implemented using the PyTorch framework. Each architecture consisted of three convolutional recurrent layers, employing either a vanilla RNN (RNN), long short-term memory (LSTM), or gated recurrent unit (GRU) cell. A BatchNorm 3D layer was applied after each of the three convolutional recurrent layers to enhance the stability and regularization of the network. To generate the predicted two-dimensional 11th frame, the output of the third layer was passed through a 2D convolutional layer. Finally, a sigmoid function was utilized to bring the pixel values within the expected range of 0 to 1, thereby producing the final predicted frame. In total, the RNN, LSTM, and GRU architectures consisted of 186 241, 3 101 377, and 556 801 parameters, respectively.

The loss function used for training was the mean squared error (MSE): 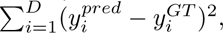 where 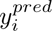 denotes the activation of pixel *i* in the 11th frame predicted by the ANN, 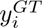 is the activation in the ground truth frame and *D* = 64 * 64 the number of pixels in the frame. Unless specified otherwise, all models were trained until overfitting was detected, which was defined as no decrease in validation loss for five consecutive epochs. The model that achieved the smallest MSE on the validation dataset, recorded five epochs prior to stopping the training, was considered the optimally trained model. This model was then used for subsequent downstream tasks, including further training on the natural video dataset in the case of the pre-trained models, thorough performance evaluation, or exploration of network characteristics. All ANNs were trained using the same set of hyperparameters to ensure comparability. To provide a reliable performance assessment accounting for potential variations in training performance, five ANNs were trained for each model. The exploration of network characteristics and performance was then based on the average results obtained from these five models.

### 4.3 Receptive field investigation

To explore the impact of pre-training on the ANN, we conducted an analysis of the receptive field properties for all training conditions. Specifically, we focused on one target neuron located at position (32, 48) in the last 2D convolutional layer, which corresponds to the layer responsible for generating the predicted 11th frame. To accomplish this, we applied a previously published activation maximization algorithm that was designed to facilitate the exploration of deep neural networks [48], to our models using a vanilla RNN architecture. By customizing the algorithm to accommodate the specifications of our network architectures, such as providing 10 input frames to the ANN, we were able to determine the input that maximized the activation of the target neuron.

The activation maximization algorithm operates by generating a random input tensor with dimensions 10 × 64 × 64, corresponding to the 10 input frames with 64 × 64 pixels each. This input tensor is then propagated through the model of interest and the resulting activation of the target neuron is obtained. Subsequently, gradient ascent is utilized to iteratively modify the input tensor, aiming to maximize the activation of the target neuron. In addition, all four regularization techniques recommended by the authors of the original algorithm were employed in this investigation to enhance the interpretability of the identified patterns [48]. Precisely, those techniques are *L*_2_ decay, Gaussian blur, and the clipping of pixels with small norm as well as those with small contribution to the activation of the target neuron. The latter results in input locations that do not contribute to the activation of the target neuron being set to zero, as those cannot be considered part of the receptive field of the respective neuron. To obtain the receptive field of the target neuron for a specific model of interest, we conducted 10 trials, each corresponding to one run of the activation maximization algorithm. This investigation was repeated for each of the 5 ANNs trained per model, resulting in a total of 5 ∗ 10 = 50 trials per model. These trials were subsequently averaged over to obtain a representative result. Within each trial, we executed 100 gradient descent steps to iteratively optimize the input tensor.

To quantify the properties of the spatio-temporal receptive fields, we explored both size and strength of the receptive field of the target neuron in each frame across different models. We identified the activated pixels (activation *>* 0.1) in each frame, counted the number of activated pixels for size assessment, and measured the 95th percentile of their activations for strength evaluation.

### 4.4 Neuronal response investigation

To further explore how pre-training shapes the neuronal response characteristics of ANNs, we analyzed neuron responses to various static grating stimuli for the following three training conditions: *DirectedProp*, a randomly initialized network, and a network fully trained on the simulated corridor dataset until detection of overfitting. As in the case of the receptive field investigation 6, we restricted our analysis to the vanilla RNN architecture. We designed the stimuli based on the Allen Brain Observatory: Visual Coding dataset [49]. The static grating stimulus consists of a full-field sinusoidal grating that varies in orientation, spatial frequency, and phase. This dataset includes 6 orientations (at 30° intervals), 5 spatial frequencies (ranging from 0.02 to 0.32 cycles per degree), and 4 phases, resulting in a total of 120 stimulus conditions.

Each grating stimulus was paired with a corresponding full-field stimulus, a uniform gray field of the same mean luminance. Both stimuli were presented for 10 frames, and the average response over this duration was computed for all neurons. For each pair, the mean response to the full-field stimulus (*F*_0_) was used as a baseline to compute the corresponding change in activity 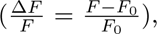 with *F* denoting the neuron’s mean response (over the 10 input frames) to the grating stimulus. This procedure yielded, for each neuron, a distribution of activity changes between each of the 120 grating stimuli and the corresponding full-field stimulus.

## Data and Code availability

All coding tasks were implemented using the Python programming language. The datasets generated and used in this study are available here: https://doi.org/10.5281/zenodo.10317798. The code used for creating, training, and evaluating the ANNs is available here: https://github.com/comp-neural-circuits/pre-training-ANNs-with-retinal-waves.

## Contributions

L.M., A.D., and J.G. designed the study; L.M. performed the research and analyzed the data related to the investigation of learning and receptive field characteristics, A.D. explored the neuronal response properties; L.M., A.D., and J.G. wrote the manuscript.

## Acknowledgements

This work was supported by the European Research Council under the European Union’s Horizon research and innovation program (NeuroDevo, Grant Agreement No. 804824) and the Technical University of Munich (TUM Innovation Network “Neurotechnology in Mental Health”). We thank Elizabeth Herbert and Shreya Lakhera for commenting on the manuscript and the entire “Computation in Neural Circuits Group” for discussions.

## Supporting information

**Fig S1. Accelerated learning in pre-trained models generalizes across various ANN architectures and retinal wave pre-training datasets. A.** Learning trajectories of four LSTM models on the simulated corridor training (left) and validation (right) datasets. Shown is the average MSE across five-fold cross-validation and the corresponding standard deviation. Training continues beyond the epochs shown, with each model stopping individually once overfitting is detected. **B.** Same as panel A, but for GRU architectures.

**Fig S2. Learning is accelerated across various pre-training dataset variations. A.** Example datapoint and learning trajectories on the propagating square dataset (Methods 4.1.3). Learning trajectories of four vanilla RNN models on the simulated corridor training (left) and validation (right) datasets. Shown is the average MSE across five-fold cross-validation and the corresponding standard deviation. Training continues beyond the epochs shown, with each model stopping individually once overfitting is detected. **B.** Same as panel A, but for the pre-training dataset consisting of retinal waves with added Gaussian blur (Methods 4.1.1). **C.** Same as panel A, but for the pre-training dataset comprising retinal waves (RWs) that propagate in all directions with a directional bias (Methods 4.1.1).

**Fig S3. Replacing training on natural stimuli with pre-training on retinal waves accelerates learning for LSTM and GRU architectures. A.** Learning trajectories of three LSTM models on the simulated corridor training (left) and validation (right) datasets. Shown is the average MSE across five-fold cross-validation and the corresponding standard deviation. Note that the performance assessment for the model pre-trained for one epoch on retinal waves (*DirectedProp one epoch* in pink) starts at epoch two, ensuring direct comparability to the *NotPretrained* model as this model does not have an advantage in training time. Training continues beyond the epochs shown, with each model stopping individually once overfitting is detected. **B.** Same as panel A, but for GRU architectures.

**Fig S4. Refinement of receptive field properties for various pre-training strategies. A.** Spatio-temporal receptive field of a target neuron in the last 2D convolutional layer of models using a vanilla RNN architecture. Shown are the 10 input frames, each cropped to a square of 27 *×* 27 pixels, which maximally activate the target neuron, indicated by a cross. **B.** Receptive field area in pixels for each frame. Note that the randomly initialized and the *DirectedProp*models are the same as shown in Fig 6. **C.** Receptive field strength, measured as the 95th percentile of pixel activations, for each frame.

