## Supplementary figures and images for "Pre-training artificial neural networks with spontaneous retinal activity improves motion prediction in natural scenes"

### Figure S1

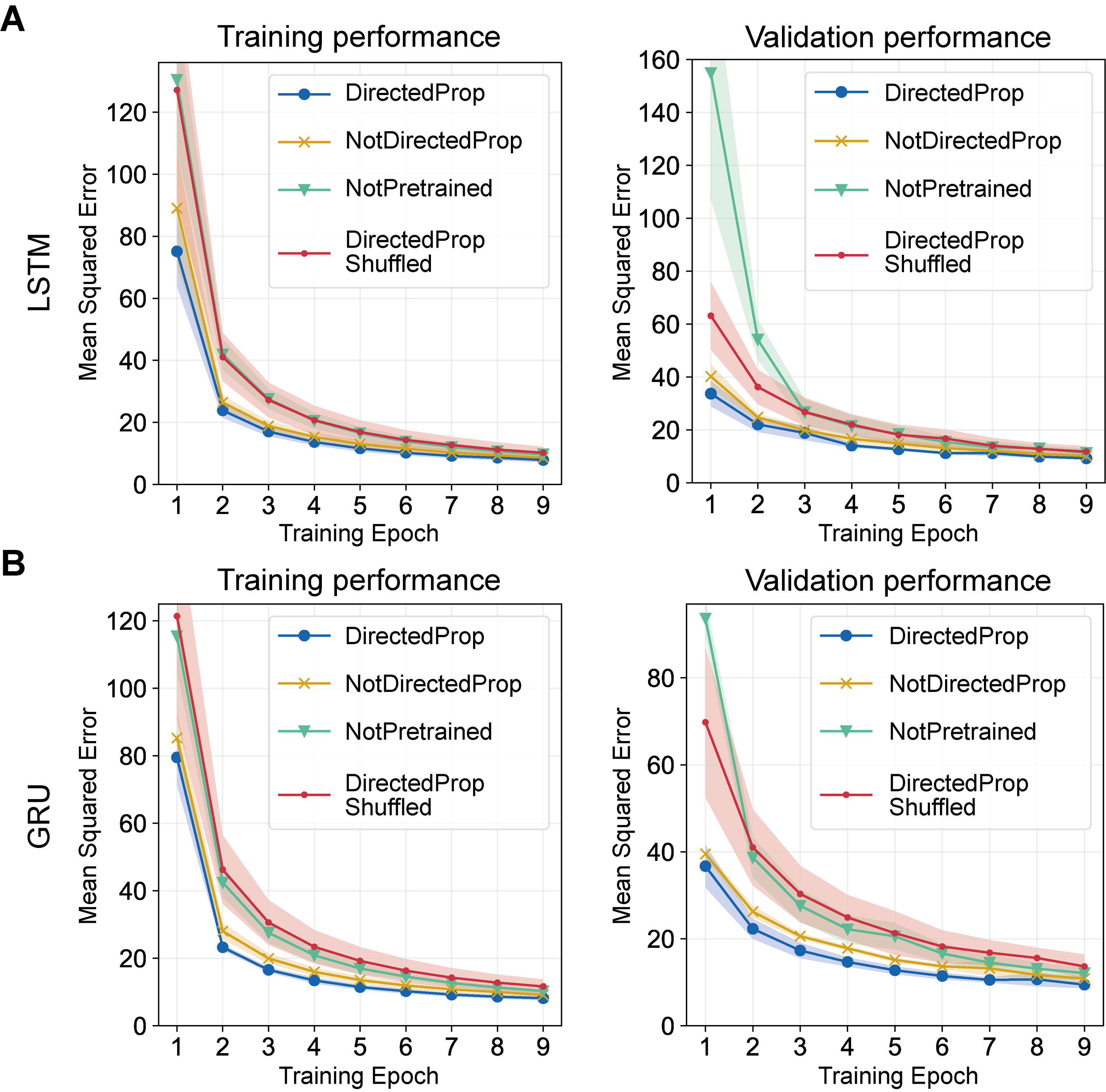

### Figure S2

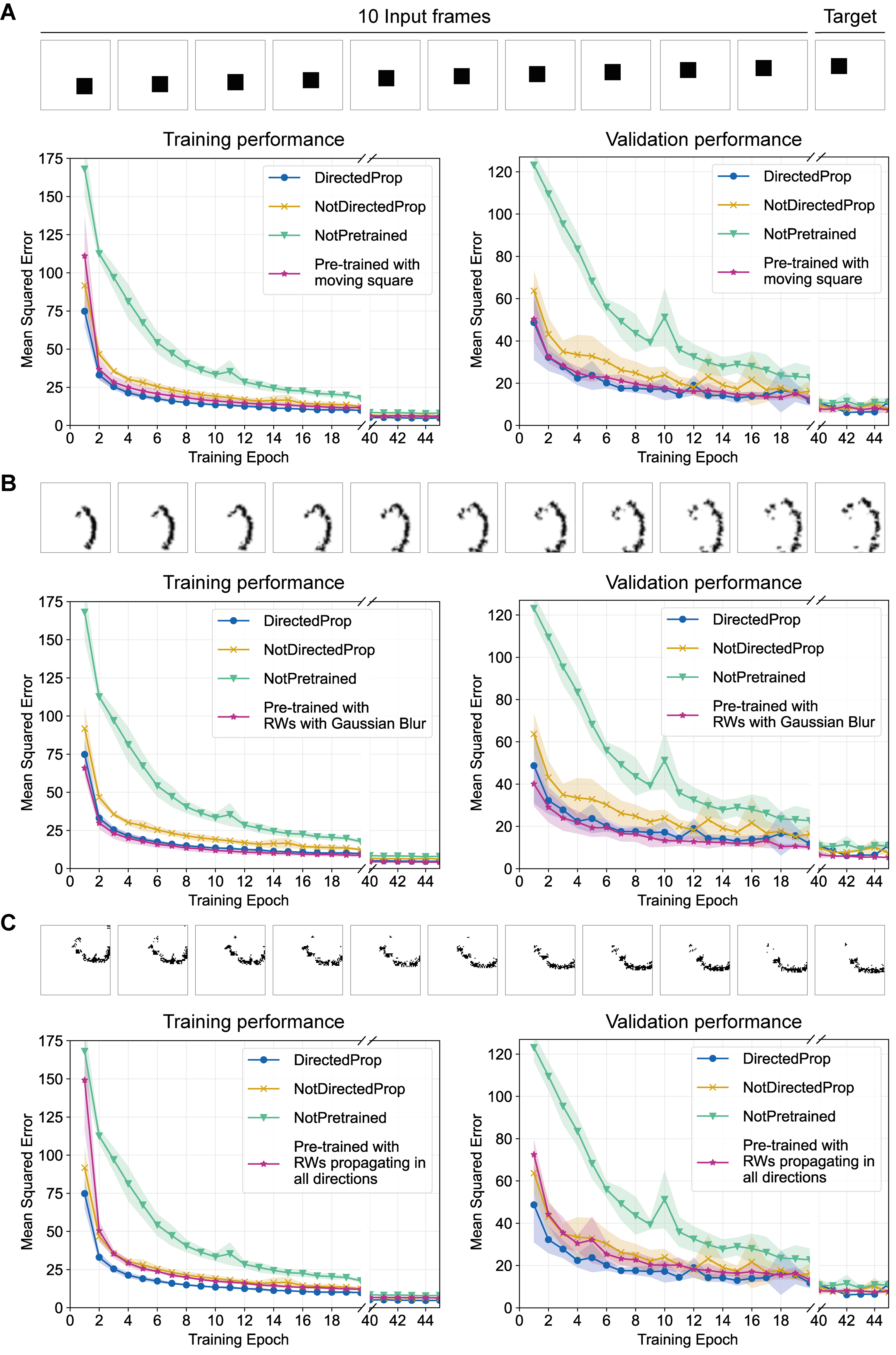

### Figure S3

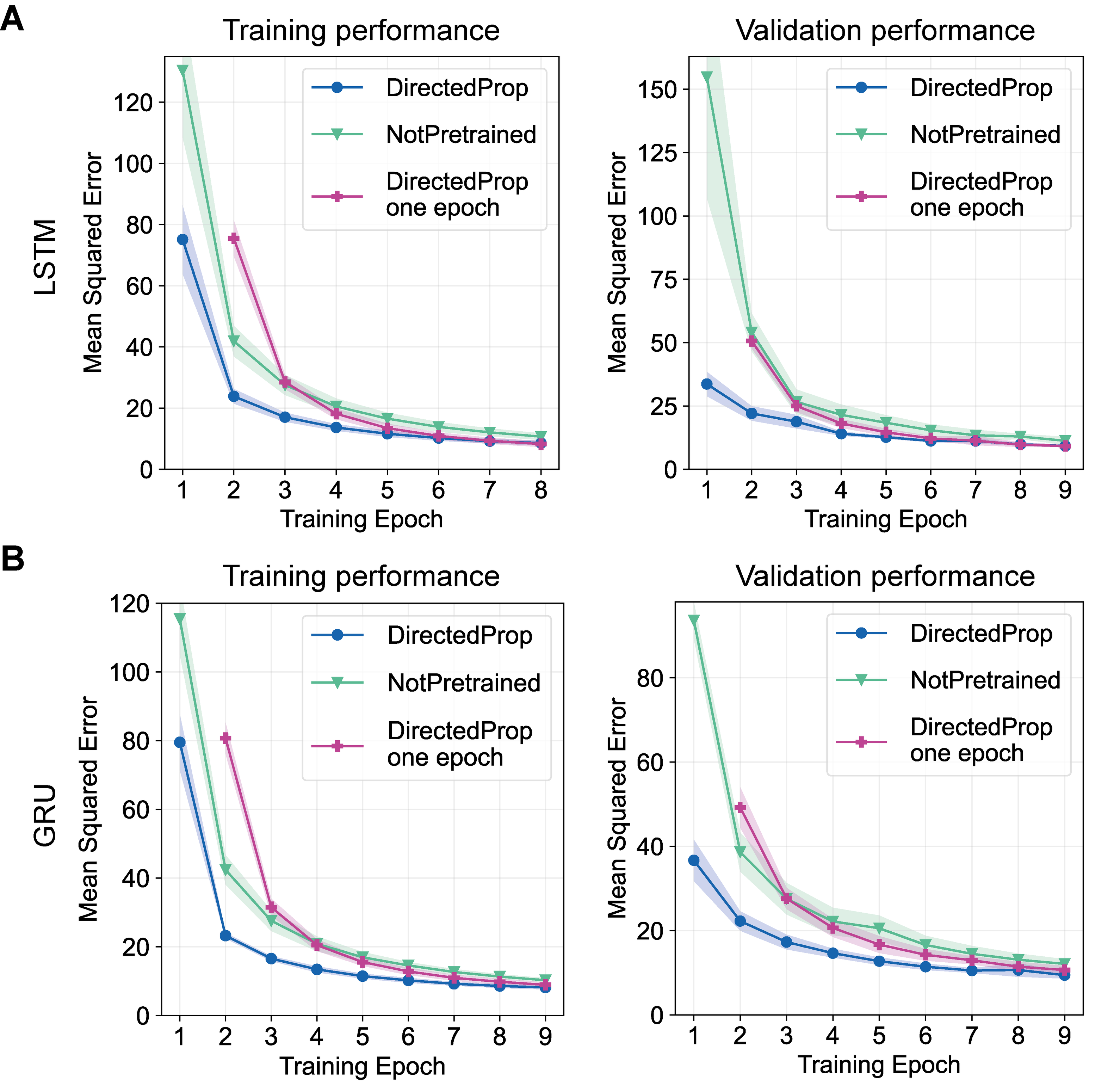

### Figure S4

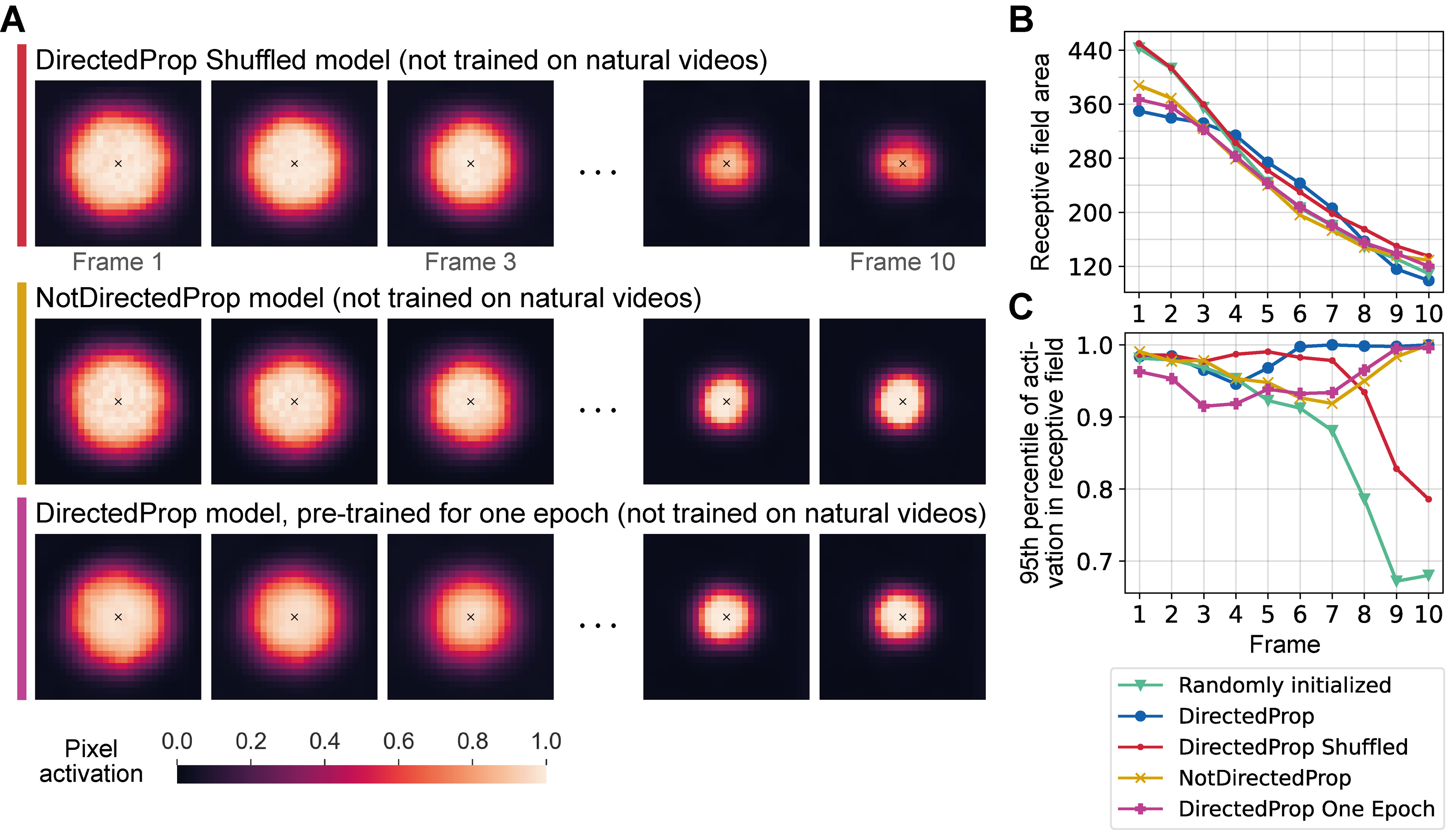
